# Epigenome-wide association study analysis of calorie restriction in humans, CALERIE™ Trial analysis

**DOI:** 10.1101/2022.06.03.494722

**Authors:** Megan E. Ramaker, David L. Corcoran, Abner T. Apsley, Michael S. Kobor, Virginia B. Kraus, William E. Kraus, David T. S. Lin, Melissa C. Orenduff, Carl F. Pieper, Reem Waziry, Kim M. Huffman, Daniel W. Belsky

**Affiliations:** Duke University Molecular Physiology Institute; Department of Genetics, University of North Carolina at Chapel Hill; The Pennsylvania State University’s Behavioral Health Department; The Pennsylvania State University’s Molecular, Cellular, and Integrative Biosciences Program; BC Children’s Hopsital Research Institute (BCCHR); Centre for Molecular Medicine and Therapeutics, University of British Columbia; Program in Child and Brain Development, CIFA, MaRS Centre; The Department of Medical Genetics, University of British Columbia; Duke University School of Medicine; Center for Aging and Human Development, Duke University Medical Center; Department of Biostatistics and Bioinformatics, Duke University Medical Center; Columbia University; Butler Columbia Aging Center, Columbia University Mailman School of Public Health; Department of Epidemiology, Columbia University Mailman School of Public Health

**Keywords:** Epigenome, Caloric Restriction, Human Aging

## Abstract

**BACKGROUND:** Calorie restriction (CR) increases healthy lifespan and is accompanied by slowing or reversal of aging-associated DNA methylation (DNAm) changes in animal models. In the Comprehensive Assessment of Long-term Effects of Reducing Intake of Energy (CALERIE™) human trial we evaluated associations of CR and changes in whole-blood DNAm.

**METHODS:** CALERIE™ randomized 220 healthy, non-obese adults in a 2:1 allocation to two years of CR or ad libitum (AL) diet. The average CR in the treatment group through 24-months of follow-up was 12%. Whole blood (baseline, 12 and 24 month) DNAm profiles were measured. Epigenome-wide association study (EWAS) analysis tested CR-induced changes from baseline to 12- and 24-months in the n=197 participants with available DNAm data.

**RESULTS:** No CpG-site-specific changes with CR reached epigenome-wide significance (FDR<0.05). Secondary analyses of CpG sites identified in published EWAS suggest, we found that CR induced DNAm changes opposite those associated with body mass index (BMI) and smoking (p<0.003 at 12- and 24-month follow-ups). In contrast, CR altered DNAm at chronological-age associated CpG sites in the direction of older age (p<0.003 at 12- and 24-month follow-ups).

**CONCLUSION:** Although individual CpG site DNAm changes in response to CR were not identified, analyses of sets CpGs identified in prior EWAS revealed CR-induced changes to blood DNAm. Altered CpG sets were enriched for insulin-production, glucose-tolerance, inflammation, and DNA-binding and -regulation pathways, several of which are known to be modified by CR. DNAm changes may contribute to CR effects on aging.

## INTRODUCTION

The geroscience hypothesis proposes that interventions that slow or reverse biological processes of aging can simultaneously prevent multiple chronic diseases and extend healthy lifespan (1). Proof of concept for geroscience is emerging from studies with animals, in which interventions that slow or reverse accumulation of molecular “hallmarks” of aging delay the onest of disease and functional impairment and extend healthy aging (2–4). One of the best-evidenced geroscience intervention in animals is calorie restriction (CR) (5). CR is defined as reduction in caloric intake from normal intake (“ad libitum” (AL)) diet while maintaining adequate nutrient intake (6). From worms to mice to monkeys, CR is associated with delayed onset of age-associated diseases, including diabetes, cancer, cardiovascular disease, osteoarthritic and increased healthy lifespan (7–10).

The mechanisms by which CR slows aging and extends healthspan in animal models are several and include alterations at physiological, metabolic, and genomic levels (6,11). Studies in animals have identified slowing or reversal of epigenetic changes associated with aging in response to CR, including alterations of whole blood DNA methylation (DNAm) (12,13). However, effects of CR on whole blood DNAm in non-obese humans are unknown.

The Comprehensive Assessment of Long term Effects of Reducing Intake of Energy (CALERIE™) study is the first long-term, randomized clinical trial of CR in healthy, non-obese humans (14). The goal of CALERIE™ was to identify the effects of 2 years of CR on predictors of longevity, disease risk factors, and quality of life. The intervention yielded substantial and sustained weight loss and signs of improved cardiometabolic health, reduced inflammation, and slowed biological aging, as measured by physiology-based algorithms (16,17). In ancillary studies in subsets of CALERIE™ participants, CR induced signs of metabolic slowing and reversal of markers of immune-system aging (18,19).

We conducted genome-wide analysis of whole-blood DNAm changes over 12 and 24 months in CALERIE™. The primary analysis tested changes in methylation levels at each of 828,613 C-G dinucleotides (CpGs). Secondary analyses tested changes at sets of CpGs identified in published epigenome-wide association studies (EWAS) of body-mass index (BMI), cigarette smoking and chronological age (20–23). BMI EWAS analysis was of interest because the CALERIE™ intervention induced substantial weight loss. Smoking- and chronological-age EWAS analyses were of interest in CALERIE™ because these are established risk factors for shortening healthy lifespan and have associations with DNAm differences at large numbers of CpG sites, which are not currently known to be directly affected by CR. We hypothesized that CR would offer a geroprotective effect which could be measured molecularly via DNAm especially at regions associated with risk factors for shorter lifespan.

## METHODS

The CALERIE™ trial randomized 220 healthy, non-obese (BMI 22.0 ≤ BMI < 28.0 kg/m^2^), adults aged 21-50 years to either a 25% calorie restriction (CR) intervention condition or ad libitum (AL) control at a 2:1 (CR:AL) ratio across three sites (Pennington Biomedical Research Center, Washington University, and Tufts University) (**Figure 1A, Table 1**) (14,24). Participants were excluded from the study if they had significant medical conditions, abnormal laboratory markers, present or potential psychiatric or behavioral problems, regular use of medications (except oral contraceptives), currently smoked, were highly physically active, or were pregnant or breastfeeding. Randomization was stratified by study site, sex and BMI. The trial duration was 24 months. As measured using doubly labeled water, the CR intervention group achieved an average of 11.7 +/− 0.7% CR (19.5+/−0.08% in the first 6 months, 9.1+/−0.7% during the subsequent 18 months) (15).

**Figure 1.**
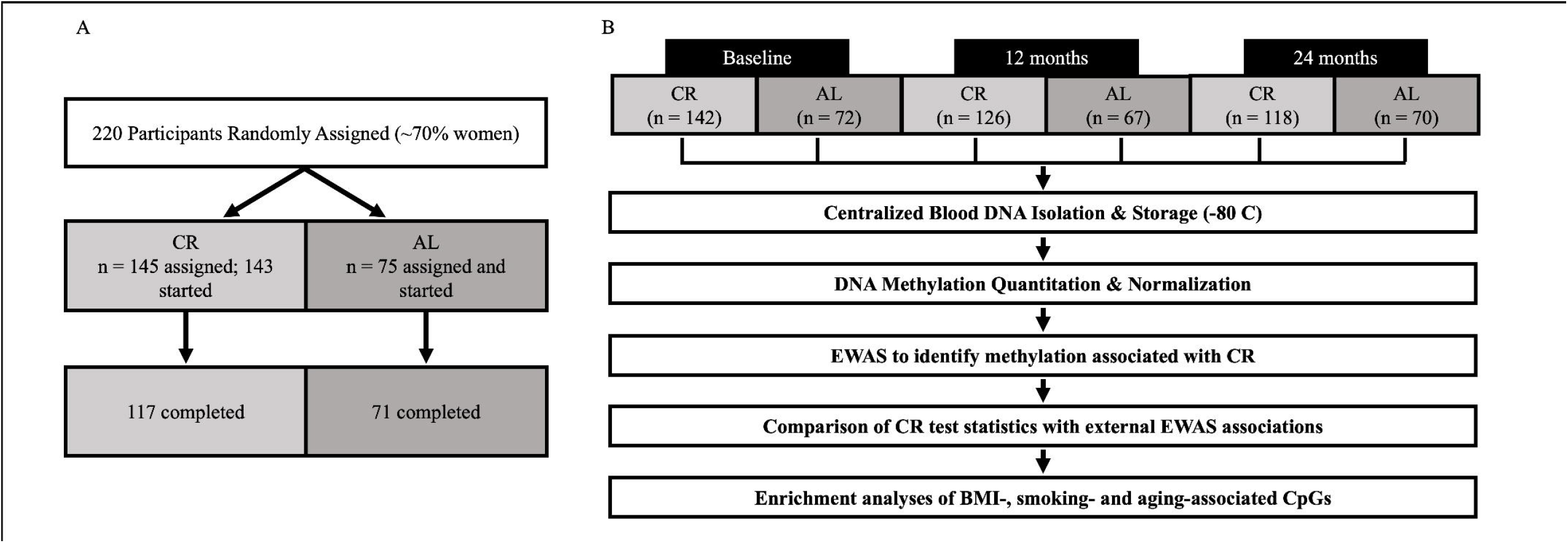
Study design. A) CALERIE trial design. 220 participants were randomly assigned to either 25% calorie restriction (CR) or ad libitum (AL) at a 2:1 ratio. Of the 220 participants assigned, 218 started and 188 completed the intervention. B) Blood samples were collected from participants at baseline and 12- and 24-month follow-ups. DNA was isolated and stored. DNA methylation was assayed with Illumina methylEPIC bead chip arrays. After quality control and normalization, epigenome-wide association study (EWAS) analysis tested CALERIE™ intervention effets at 12- and 24-month follow-ups at each of 828,613 CpG sites. Finally, we conducted secondary analysis comparing results from CALERIE EWAS with results from published EWAS of BMI, cigarette smoking, and chronological age.

**Table 1.**
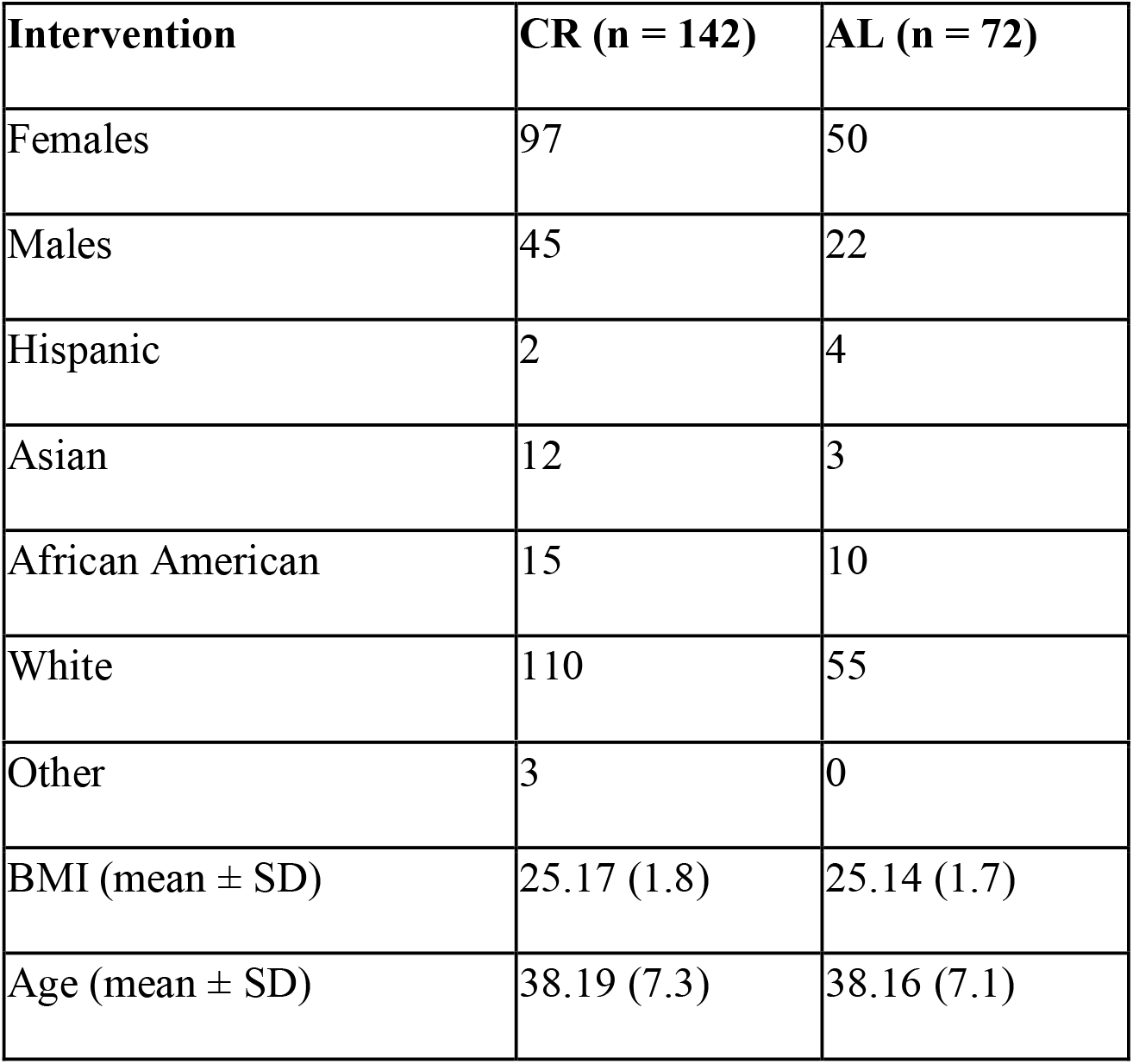
Study participant characteristics at pre-treatment baseline.

### DNA Methylation

DNAm profiling was conducted in the Kobor Lab from whole-blood DNA stored at −80°C. After quality controls and normalization, DNAm datasets were generated for n=595 samples from 214 individuals (142 CR, 72 AL) (**Figure 1B**, **Table 1**). Briefly, 750 ng of DNA was extracted from whole blood and bisulfite converted using the EZ DNA Methylation kit (Zymo Research, Irvine, CA, USA). Methylation was measured from 160 ng of bisulfite-converted DNA using the Illumina EPIC Beadchip (Illumina Inc, San Diego, CA, USA). Quality control and normalization were performed using methylumi (v. 2.32.0) (25) and the Bioconductor (v 2.46.0) (26) packages from the R statistical programming environment (v 3.6.3). Probes with detection p-values >0.05 were coded as missing; probes missing in >5% of samples were removed from the dataset (final probe n=828,613 CpGs). Normalization to eliminate systematic dye bias in the the 2-channel probes was carried out using the methylumi default method. We conducted principal component analysis of EPIC-array control-probe beta values to compute controls for technical variability across the samples (27).

### Statistical Analysis

Primary analysis was an epigenome-wide association study (EWAS) of CALERIE™ treatment effects in which treatment group was the exposure and changes in probe beta value from baseline to 12 months and baseline to 24 months were the outcome variables.

Secondary analyses examined sets of CpG sites identified in published EWAS of obesity, cigarette smoking and chronological age to test if CALERIE™ treatment specifically affected DNAm at CpG sites known to be altered by these exposures.

#### Epigenome-wide Association Study (EWAS) of CALERIE™ Treatment Effects

We tested associations of CALERIE™ intervention with changes in DNAm at each QC’ed CpG site using a mixed model. The model took the form of:

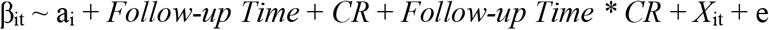

where ‘β’ is the level of methylation for CpG site ‘i’ at time ‘t’; ‘a’ is the model intercept, including sample-wide and person-specific components, ‘*Follow-up Time*’ is a pair of indicator variables encoding the 12- and 24-month follow-ups; ‘*CR*’ is an indicator of treatment group; *‘Follow-up Time * CR’* is a series of interaction terms between follow-up time and treatment group; *‘X’* is a matrix of covariates; and ‘e’ is the error term comprising both sample-wide and person-specific components. The effect of intervention is tested by the coefficients for the interaction terms, which evaluate the treatment effect at 12 and 24 months as the difference in change from baseline between the treatment (CR) and control ad-libitum (AL) groups.

Time-invariant covariates were pre-intervention-baseline chronological age, sex, BMI stratum (22-24.9, 25-27.9), study site, and the first three principal components estimated from genome-wide SNP data. Time-varying covariates were proportions of monoctyes, neutrophils, and CD4T, CD8T, Natural Killer, and B-cell lymphocytes estimated from the DNAm data using the Houseman Equation via the Minfi and FlowSorted.Blood.EPIC R packages and the first seven principal components estimated from EPIC-array control probes (27–29). Benjamini-Hochberg correction was applied to account for non-independence of tests. Statistical significance was established at a false discovery rate (FDR) <0.05. EWAS analysis was conducted using the lmerTest R package (30).

#### Secondary Analyses of EWAS Summary Statistics

We evaluated whether DNAm changes associated with CALERIE™ intervention reflected changes expected based on published EWAS. We conducted analyses of EWAS results from studies of body-mass index (BMI), cigarette smoking, and chronological age (20–23). Hypothesis testing was performed using a Wilcoxon Rank-Sum test to compared distributions of CALERIE™ EWAS test statistics for phenotype-associated CpGs to the distribution of CALERIE™ EWAS test statistics for all other CpGs. Independent tests were performed for CpG sites identified as hypermethylated and hypomethylated in association with the target phenotype. Because all target-phenotype EWAS used an earlier generation of Illumina array technology, we restricted this analyses to the 431,205 EPIC-array CpGs measured in CALERIE™ that were also included on the Illumina 450k array.

##### Secondary analysis of BMI-associated CpGs

The CALERIE™ intervention was associated with an average weight loss of 8 kg by 12 months of follow-up (15). We therefore evaluated whether DNAm changes associated with the CALERIE™ intervention overlapped with DNAm associations with BMI. We examined 129 CpGs identified in a prior EWAS of BMI (20). Specifically, we tested if CpGs hypomethylated in individuals with higher BMI showed signs of increased DNAm in response to the CALERIE™ intervention, and if CpGs hypermethylated in individuals with higher BMI showed signs of decreased DNAm in response to CALERIE™ intervention; i.e., we tested the hypothesis that DNAm changed induced by CALERIE™ intervention would be opposite to the pattern of association with higher BMI.

##### Secondary analysis of smoking-associated CpGs

We tested if DNAm changes associated with the CALERIE™ intervention overlapped with DNAm associations with cigarette smoking, a potent risk factor for aging-related disease and mortality known to have pervasive effects on blood DNAm. We examined 2,622 CpGs identified in a prior EWAS of smoking (23). We tested if CpGs hypomethylated in smokers showed signs of increased DNAm in response to the CALERIE™ intervention and if CpGs hypermethylated in smokers showed signs of decreased DNAm in response to the CALERIE™ intervention; i.e. we tested the hypothesis that DNAm changes induced by CALERIE™ intervention would be opposite to the pattern of association with smoking.

##### Secondary analysis of chronological-age-associated CpGs

We tested if DNAm changes associated with the CALERIE™ intervention overlapped with DNAm associations with chronological age. We examined 1,000 CpGs identified in a prior EWAS of chronological age (21). We tested if CpGs hypomethylated in chronologically older individuals showed signs of increased DNAm in response to the CALERIE™ intervention and if CpGs hypermethylated in chronologically older individuals showed signs of decreased DNAm in response to CALERIE™ intervention; i.e. we tested the hypothesis that DNAm changes induced by CALERIE™ intervention would be opposite to the pattern of association with older chronological age. We repeated the analysis using 875 CpGs identified in a prior EWAS of chronological age (22).

For all secondary analysis, we applied a Bonferoni-corrected threshold of p<0.003 to establish statistical significance (16 tests; 0.05/16=0.003).

### Enrichment Analyses

To inform interpretation of secondary analyses, we performed enrichment analysis of sets of CpGs identified in published EWAS (20–22). We annotated each CpG to the nearest transcription start site (TSS) to conduct gene enrichment analysis. We used the Reactome Database to identify enriched biological processes and functional relationships (31). We used the GM12878 chromatin immunoprecipitation sequencing (ChIP-seq) data from the ENCODE data portal (32) to identify whether certain transcription factor binding sites (TFBSs) were enriched amongst phenotype-associated CpGs. Briefly, BEDtools was used to identify the intersection between the Methyl450 annotation file and the ChIP-seq bed file (33). Enrichment of transcription factors bound within 500 bp of the phenotype-associateed CpGs compared to non-phenotype-associated CpGs was tested with permutation analysis. We tested ontological enrichment using the gene ontology enrichment analysis and visualization tool (Gorilla) (34).

## RESULTS

### EWAS of CALERIE™ Treatment Effects

We conducted intent-to-treat (ITT) analysis of CALERIE™ treatment effects at 12- and 24-month follow-ups. Genome-wide comparison of DNAm between CR and AL at 12 and 24 months did not identify any CpG-site-specific changes that were statistically different from zero at FDR<0.05 (**Figure 2**; **Supplementary Table 1**). The top-ranked CpG site at 12 months was within the first exon of T-Cell Receptor T3 Delta Chain (CD3D) (cg07728874, p-value = 4.05×10^-6^). At 24 months, the top-ranking CR-associated site was located on chromosome 1 within Long intergenic Non-Protein Coding RNA 1344 (LNC01334) (cg12040931, p-value = 2.5×10^-6^).

**Figure 2.**
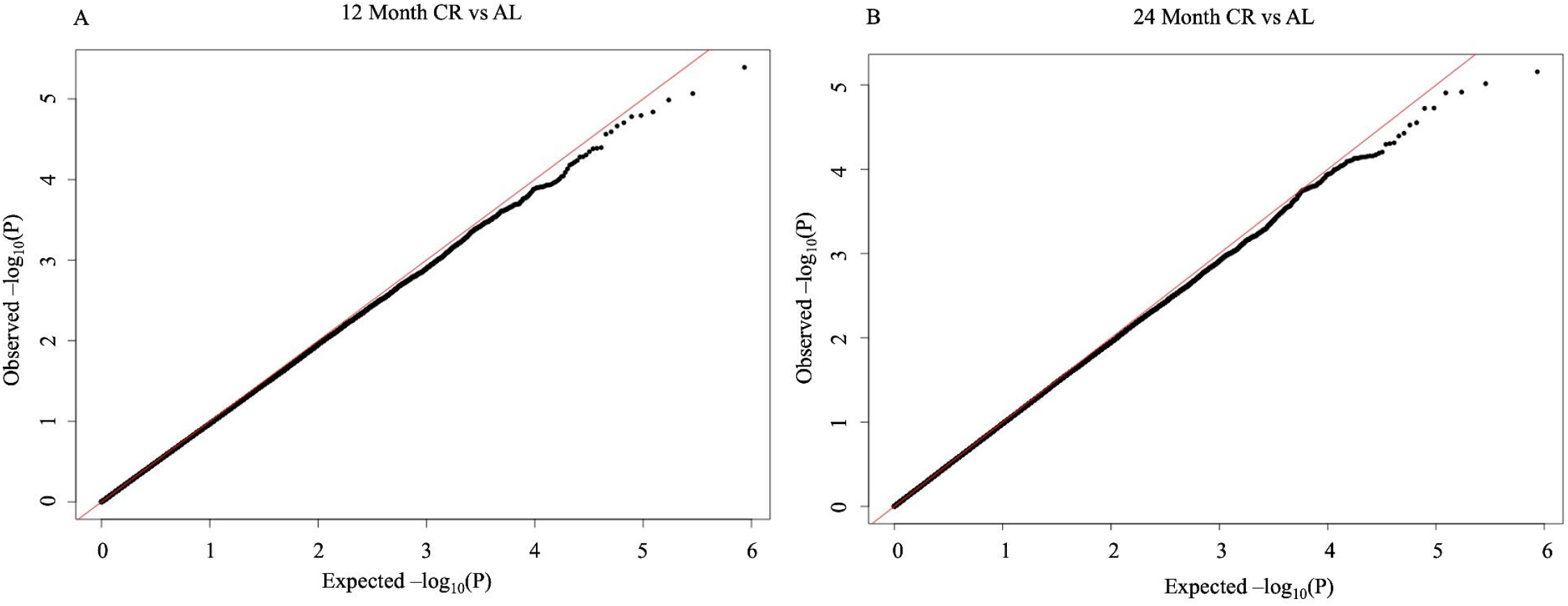
Quantile-quantile (QQ) plots of p-value distributions from epigenome-wide association study (EWAS) analysis of CALERIE treatment effects at 12- and 24-month follow-ups. The figure shows QQ plots for EWAS of blood DNA methylation changes in response to CR at 12-months (Genomic Inflation – 0.97; Panel A) and 24-months (Genomic Inflation – 0.99; Panel B).

### Secondary Analyses of CpG sites identified in published EWAS of BMI, cigarette smoking, and chronological age

We conducted secondary analyses of summary statistics from the CALERIE™ EWAS using published results from EWAS of BMI, cigarette smoking and chronological age. CR-induced substantial weight loss (15).

We first compared CALERIE™ EWAS results for DNAm at n=129 CpG sites identified in a published EWAS of BMI (20) with results for all other CpG sites. For CpG sites identified as hypermethylated in individuals with higher BMI (n=50), CR tended to reduce DNAm (12-month p=2.06E-07; 24-month p=3.96E-11). For CpG sites identified as hypomethylated in individuals with higher BMI (n=79), CR tended to increase DNAm (12-month p=1.04E-06; 24-month p=7.04E-04). Thus, for both sets of CpGs, CR reversed BMI-associated DNAm.

We next compared CALERIE™ EWAS results for DNAm at n=2,622 CpG sites identified in EWAS of cigarette smoking(35) with results for all other CpG sites. For CpG sites identified as hypermethylated in smokers (n=1,555), compared with AL, CR tended to reduce DNAm (12-month p=1.03E-05; 24-month p=2.63E-30). For CpG sites identified as hypomethylated in smokers (n=1,067), compared with AL, CR tended to increase DNAm, although this finding was statistically different from the null only at 24 months of follow-up (12-month p=0.08; 24-month p=4.3E-04). Overall, CR showed signs of reversing smoking-associated DNAm.

Finally, we compared CALERIE™ EWAS results for DNAm at 1,000 CpG sites previously associated with chronological age (21) to results for all other CpG sites. For CpG sites identified as hypermethylated in older adults (n= 980), compared with AL, CR tended to increase DNAm (12-month p=3.79E-41; 24-month p=5.73E-06). For CpG sites identified as hypomethylated in older adults (n=20), compared with AL, CR was not associated with changes in DNAm (12-month p=0.12; 24-month p= 0.29). Results were similar in repeated analyses using results from a second EWAS of chronological age (22). Thus, for sites hypermethylated in older adults, CR induced DNAm changes consistent with older age. In contrast, CR had no detectable effect on sites hypomethylated in older as compared to younger adults.

Results for analyses of BMI-, cigarette smoking-, and chronological-age-associated CpG sites are reported in **Table 2**. Distributions of CALERIE™ EWAS test statistics for BMI-, cigarette-smoking, and chronological age-associated CpGs are shown in **Figure 3**. Enrichment results are reported in **Supplementary Table 2.** External EWAS CpGs and test statistics are included in **Supplementary Table 3**.

**Figure 3:**
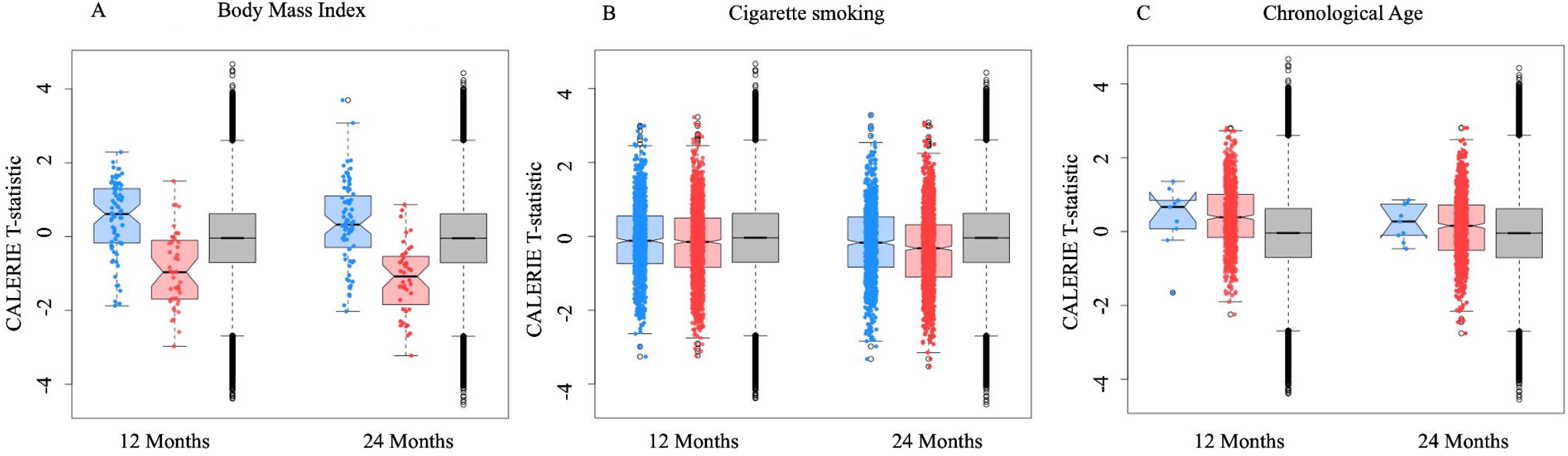
Distributions of test statistics from epigenome-wide association study (EWAS) analysis of CALERIE™ treatment effects for CpG sites identified in published EWAS of body mass index, cigarette smoking, and chronological age. The figure shows box plots of CALERIE™ EWAS test statistics for three groups of CpGs sites for each target phenotype. The blue-shaded box plot shows CALERIE™ EWAS test statistics for CpG sites that exhibit lower levels of DNA methylation in association with the target phenotype in published EWAS of independent samples. The red-shaded box plot shows CALERIE™ EWAS test statistics for CpG sites that exhibit higher levels of DNA methylation in association with the target phenotype in published EWAS. The gray-shaded box plot shows test statistics for CpG sites not associated with the target phenotype in published EWAS. Box plots are drawn for CALERIE™ EWAS results from 12- and 24-month follow-ups. Stars indicate p-value threshold for comparisons based on a Wilcoxon Rank Sum Test. (* < 0.05, ** < 0.005, *** < 0.0005). Panel A graphs data grouped according to EWAS of Body Mass Index by Wahl et al. (2017). The figure illustrates reversal of BMI-associated DNAm changes in response to CALERIE™ intervention. Panel B graphs data grouped according to EWAS of cigarette smoking by Joehanes et al. (2020). The figure illustrates reversal of smoking-associated DNAm changes in response to CALERIE™ intervention. Panel C graphs data grouped according to EWAS of chronological age by McCartney et al. 2020. The figure illustrates induction of older-chronological-age-associated DNAm changes in response to CALERIE™ intervention, although only for sites exhibiting increased DNAm in older as compared to younger people.

**Table 2.**
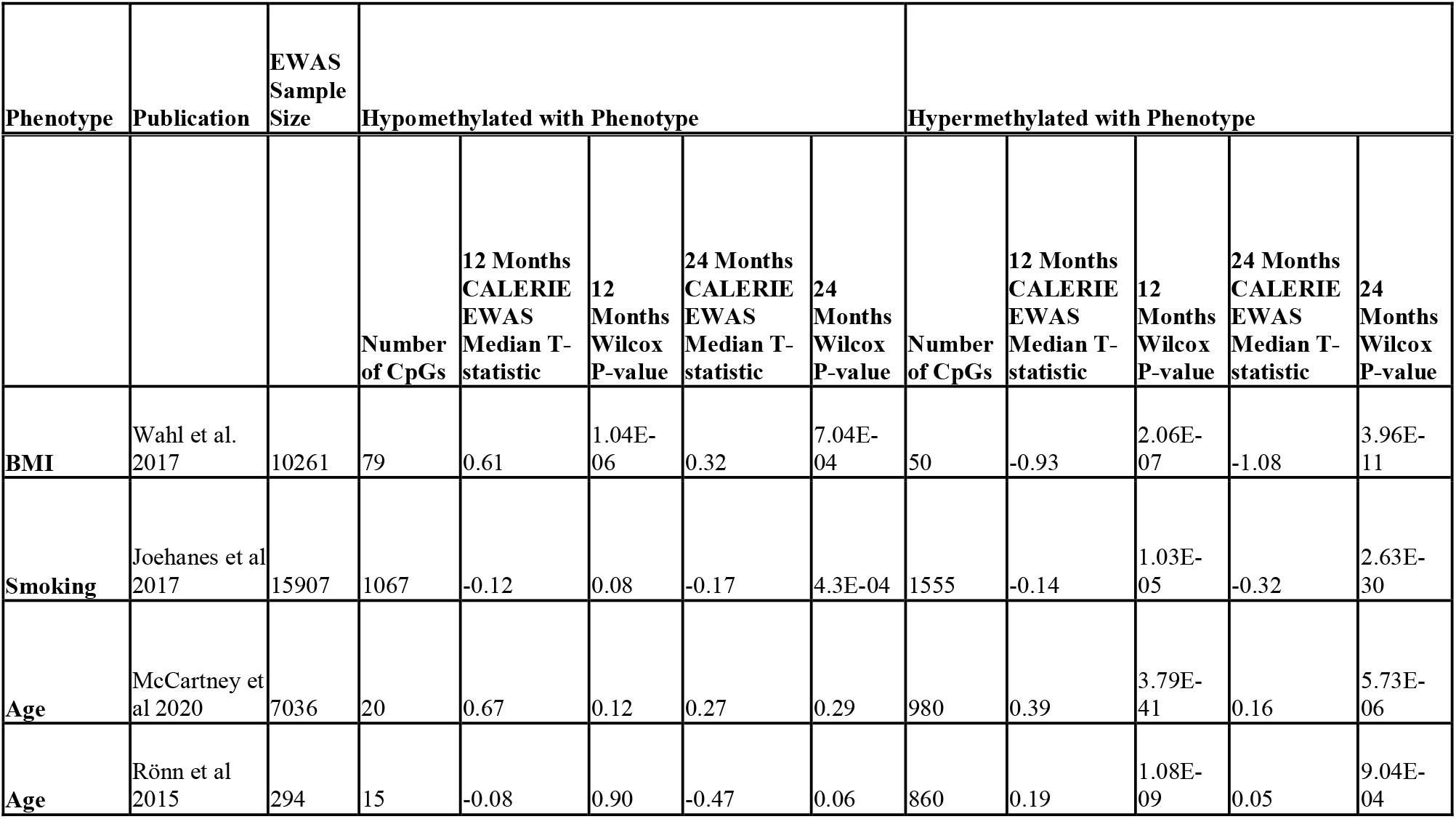
Analysis of CpG sites identified in published epigenome-wide association studies of body-mass index, smoking, and chronological age. The table summarizes results for anlaysis of CpG sites identified in published epigenome-wide association studies (EWAS) of body-mass index (BMI), smoking, and chronological age. We included CpGs identified in the published EWAS with p-value<1E-7, with the exception of the chronological-age EWAS by McCartney et al., which reported only the top 1000 sites (all p-values<1E-7). We used a Wilcoxon Rank-Sum to compare CALERIE-EWAS T-statistic distributions between phenotype-associated CpGs and all other CpGs. Tests were conducted separately for CpGs hypermethylated with the phenotype (i.e. for which the association between the phenotype and DNA methylation was positive) and CpGs hypomethylated with the phenotype (i.e. for which the association between the phenotype and DNA methylation was negative). The table reports the number of CpGs included in each test and the resulting z-statistic and p-value. Positive median CALERIE EWAS t-statistics indicate that DNAm change in response to CALERIE intervention was in the same direction as the DNAm association with the target phenotype. A negative median t-statistic indicates that the DNAm change in response to CALERIE intervention was in the opposite direction of DNAm association with the target phenotype.

## DISCUSSION

The goal of the CALERIE™ Trial was to identify effects of CR on predictors of longevity, disease risk factors and quality of life. Published analyses of CALERIE™ data establish that the intervention improved cardiometabolic health and suggest it may have slowed or reversed aging-related biological changes (15–19,36). In this study, we tested whether the intervention altered whole-blood DNAm. After accounting for multiple testing, EWAS analysis revealed no sites of altered CpG methylation by CR. However, secondary analyses of sets of CpG sites, identified in published EWAS of BMI, cigarette smoking, and chronological age, indicated that the CALERIE™ intervention changed blood DNAm in a manner consistent with a reversal of DNAm patterns linked with obesity and cigarette smoking, but in the direction of older chronological age. Further interrogation across BMI-, cigarette smoking-, and chronological aging-associated sites revealed enrichment for pathways involved in insulin production, glucose tolerance, inflammation, and DNA binding and regulation (**Supplementary Table 2**).

CALERIE™-induced DNAm changes at BMI-associated CpG sites were enriched for genes involved in insulin production, glucose tolerance and inflammatory processes, consistent with CR-induced epigenetic changes in animal models (7–10,32,37–40). The 26 genes enriched in CpG sites hypermethylated with higher BMI include P4HB, critical for lipoprotein metabolism, insulin production, and glucose intolerance (37–39). CR induced hypomethylation at P4HB may mediate previously reported CR-derived metabolic improvements in lipoproteins and insulin sensitivity (17). Another potential epigenetic benefit of CR on glucose tolerance may derive from hypermethylation at cg16246545 (**Supplementary Table 4**), located near PHGDH. Deletion of PHGDH in adipocytes of mice with diet-induced obesity improves glucose tolerance. CR-induced methylation changes at both P4HB and PHGDH likely enhance glucose tolerance. Additional CR-induced epigenetic changes at BMI-associated sites included hypomethylation at cg19750657 (**Supplementary Table 4**), located near UFM1, which has been identified as a mediator of the inflammatory response in diabetic mice. Taken together, these results imply that CR, especially when maintained for 24 months, may produce anti-inflammatory benefits (32,40).

CALERIE™-induced DNAm changes at smoking-associated CpG sites were enriched for genes involved in the tumor necrosis factor receptor-2 (TNF2) non-canonical NF-kB signalling pathway, a key driver of systemic inflammation (41). In addition, changes at sites with less methylation in smokers vs. non-smokers included sites identified in published EWAS of C-reactive protein (CRP) (42), a well-studied biomarker of inflammation, which is elevated in smokers and was reduced with CR in CALERIE™ (43–47). Taken together, CR appears to reverse smoking-associated DNAm patterns in inflammatory pathways.

The overwhelming majority of CpG sites identified in EWAS of chronological age exhibited greater DNAm in older as compared to younger individuals. These sites, at which we observed increased DNAm in response to CR, are enriched for multiple transcription factors and DNA binding proteins including T-Box Transcription Factor 15 (TBX15), SRY-Box Transcription Factor 1 (SOX1), Zic Family Member 4 (ZIC4), SIM BHLH Transcription Factor 1 (SIM1), and SRY-Box Transcription Factor 17 (SOX17). Therefore, CR may induce gain of methylation parallel to aging at genomic sites serving regulatory functions. An important next step is to better understand if such gain of methylation reflects processes of aging-related decline in system integrity or, instead, genomic changes that preserve health in aging. For example, the association of CpG methylation at these sites with chronological age could reflect survivor bias, in which relatively fewer individuals with lower levels of DNAm at these sites survive to advanced ages. CR slows the accumulation of aging-related DNAm changes in mice and monkeys (12,13). Further investigation of the significance of chronological-age-associated CpG sites for phenotypes of aging is needed to clarify interpretation of our findings.

We acknowledge limitations. Foremeost, response to the CR intervention was heterogeneous, as is typical in lifestyle interventions (49). Over the 2-year intervention, the treatment group achieved on average 12% CR (15). The trial sample was relatively small for genome-wide analysis; EWAS analyses were powered to detect only medium-to-large effect-size changes in DNAm at individual CpG sites. Identification of such changes is hampered by imperfect measurement precision for individual CpG-site DNAm (49), which will bias estimates of change toward the null. Nevertheless, aggregate analyses of sets of CpGs identified in prior EWAS suggest that the CALERIE™ intervention altered the blood methylome. As new methods are developed to improve precision of DNAm measurement from Illumina array data, it may be possible to revisit analyses to identify specific regions in which DNAm may be altered by the intervention (50). Future studies testing stronger doses of CR or including larger samples may also improve detection of DNAm changes. Last, because follow-up extended only to the end of the intervention period, we cannot know if DNAm changes associated with CR persisted after the intervention concluded.

In conclusion, while CR did not result in individual CpG-site DNAm changes that reached epigenome-wide significance, analyses of sets of CpGs identified in prior EWAS of BMI, cigarette smoking and chronological age identified clear evidence of DNAm changes in response to CR. As expected, the BMI-associated changes were consistent with CR-induced reversal of a BMI-associated patterns of DNAm. Likewise, CR reversed DNAm patterns associated with cigarette smoking, a known correlate of premature aging. Last, and to our surprise, CR appeared to increase methylation at sites where hypermethylation is associated with older as compared to younger age. That these sites were enriched for regulatory mechanisms suggests complex interplay of CR with genomic changes characteristic of older age. Whether they imply pro-aging effects of CR or reflect signatures of healthy aging remains to be determined.

## Supporting information

Enrichment Analyses

## Acknowledgement

This work was supported by National Institute on Aging Grant R01AG061378. WEK and KMH are also supported by the National Institute of Aging Grant R33AG070455. VBK is supported by National Institute of Health Grants R01AG054840 and P30-AG028716. RW is the recipient of a McKnight Scholar Award from the McKnight Brain Research Foundation through the American Brain Foundation in collaboration with the American Academy of Neurology.

## Conflicts of Interest

None

We thank the CALERIE Biorepository (R33AG070455) for its support of this work.

## Notes

### Competing Interest Statement

The authors have declared no competing interest.

