## Supplementary material for "Epigenome-wide association study analysis of calorie restriction in humans, CALERIE™ Trial analysis": Enrichment Analyses

**Supplemental Methods to Epigenome-wide association study analysis of calorie restriction in humans, CALERIE^TM^ Trial analysis**

We performed enrichment analyses amongst previously defined BMI- and aging-associated CpGs to further interpret the biological significance of altered methylation at those sites in response to CR (1–3). Gene enrichment was conducted by annotating each CpG site to the nearest TSS and performing permutation testing to identify whether certain genes were enriched amongst BMI- or aging-associated CpGs. The Reactome Database was utilized to identify enriched biological processes and functional relationships (4). To identify whether certain transcription factor binding sites (TFBSs) were enriched amongst phenotype-associated CpGs the GM12878 chromatin immunoprecipitation sequencing (ChIP-seq) data from the ENCODE data portal was used (5). Briefly, BEDtools was used to identify the intersection between the Methyl450 annotation file and the ChIP-seq bed file (6). Permutation testing then allowed for the identification of whether specific transcription factors bound within 500 bp of the phenotype-associated CpGs were enriched compared to non-phenotype-associated CpGs. Furthermore, ontological enrichment of genes nearest hyper- and hypomethylated BMI- and aging-associated CpGs was performed using the gene ontology enrichment analysis and visualization tool (Gorilla) (7).

*Enrichment in BMI-, smoking- and chronological-age-associated sites*. Of the sites hypomethylated with greater BMI (n = 79), 11 genes and 1 transcription factor (NF-Kappa-B Transcription Factor P65 (RELA)) were enriched (Supplementary Table 2, Supplementary Figure 1) (3). Of the sites hypermethylated with greater BMI (n = 50), 26 genes were enriched; no transcription factor binding sites were enriched (Supplementary Table 2 Supplementary Figure 1). There were no pathways enriched in either BMI- or aging-associated CpG sets regardless of methylation association. The most enriched ontological processes in hypermethylated BMI-associated sites included processes involved in regulation of protein transport and localization. The most enriched ontological processes in the set of hypomethylated BMI-associated sites included processes involved in carboxylic acid and amino acid transport as well as regulation of lipid kinase activity.

Of the combined set of CpG sites hypomethylated with older age (n = 35) from the two EWAS studies used for comparison to the CALERIE results, 18 genes were enriched and 1 transcription factor (EBF1) had enriched binding (Supplementary Table 2, Supplementary Figure 2) (1,2). Of the combined set of CpG sites hypermethylated with older age (n = 1,741), 108 genes were enriched and 2 transcription factors (Enhancer of Zeste 2 Polycomb Repressive Complex 2 Subunit, EZH2, and RE1 Silencing Transcription Factor, REST) had enriched binding (Supplementary Table 2, Supplementary Figure 2). The most enriched ontological process in sites that have more methylation with older age was “response to purine-containing compound.” Additionally, enriched ontological functions amongst the same sites included DNA-binding transcription factor activity and regulatory region DNA binding.

**Supplementary Table 1. Epigenome-wide Association Study (EWAS) results for tests of CALERIE treatment effects at 12- and 24-month follow-ups.** The table reports results for each CpG site included in analysis. Coefficient estimates reflect the estimated difference between the CR treatment and AL control groups in the within-individual change in DNA methylation beta values relative to baseline. [then copy paste the methods section test describing the EWAS models]. Columns in the EWAS results table are named as follows [list each column header and explain what it means. If you prefer, you can include a mini table here that lists the column headers in the left column and definitions of what they correspond to in the right column]

Complete EWAS results are available upon request.

**Supplementary Table 2. EWAS enrichment results.**

**Supplementary Table 3. External EWAS CpGs and test statistics.**

**Supplementary Table 4. BMI-associated CpGs of interest.**

**Supplemental Figure 1.** Flow diagram of enrichment analyses performed on BMI-associated CpGs from Wahl et al 2017. CpGs were divided into those that have increased methylation with higher BMI (n = 50) and those that have decreased methylation with higher BMI (n = 79). Gene, pathway, and transcription factor binding site (TFBS) enrichment analyses were performed. The number of enrichments in each category, colored by relationship of methylation with chronological age (red = increased methylation, blue = decreased methylation) are indicated in the bottom of the flow diagram.

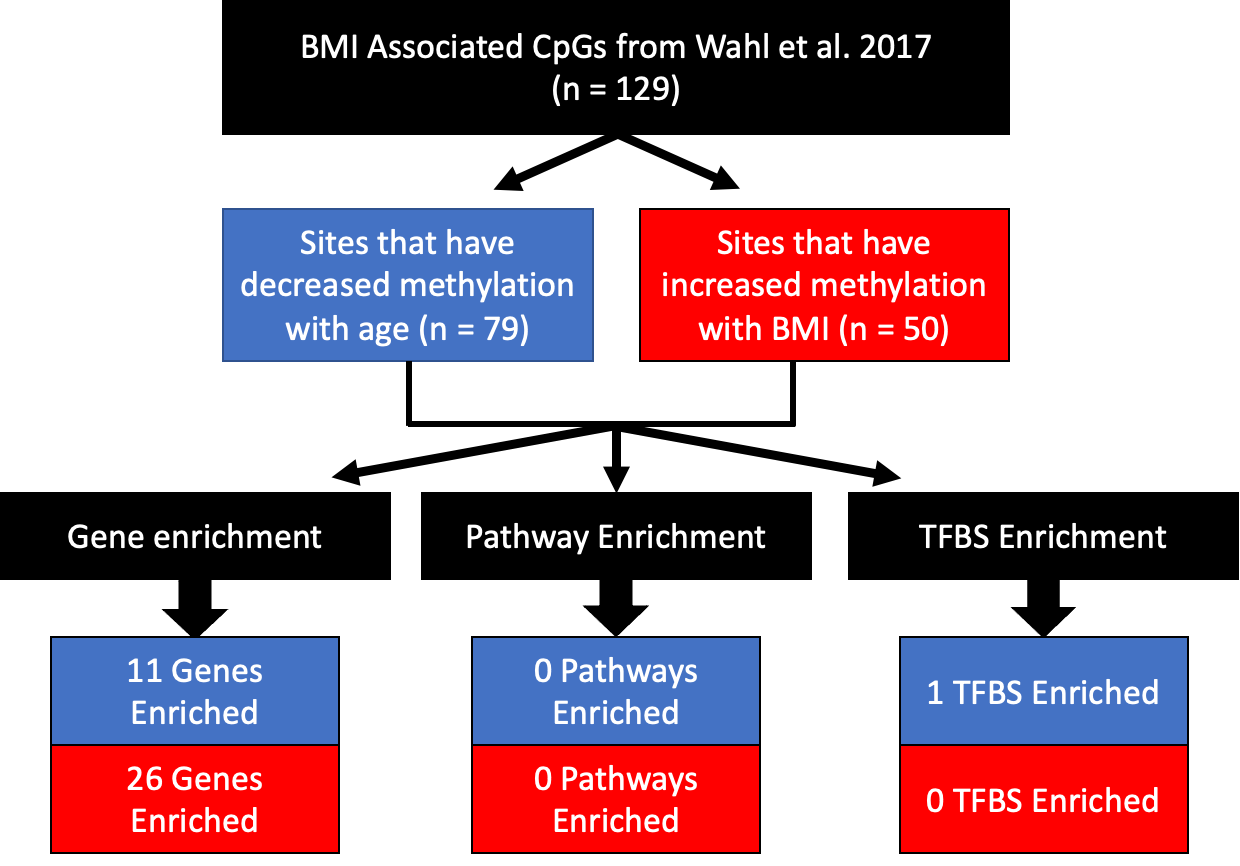

**Supplemental Figure 2.** Flow diagram of enrichment analyses performed on chronological age associated CpGs from both McCartney et al. 2020 and Ronn et al. 2015. CpGs were divided into those that have increased methylation with age (n = 1741) and those that have decreased methylation with age (n = 35). Gene, pathway and transcription factor binding site (TFBS) enrichment analyses were performed. The number of enrichments in each category, colored by relationship of methylation with chronological age (red = increased methylation, blue = decreased methylation) are indicated in the bottom of the flow diagram.

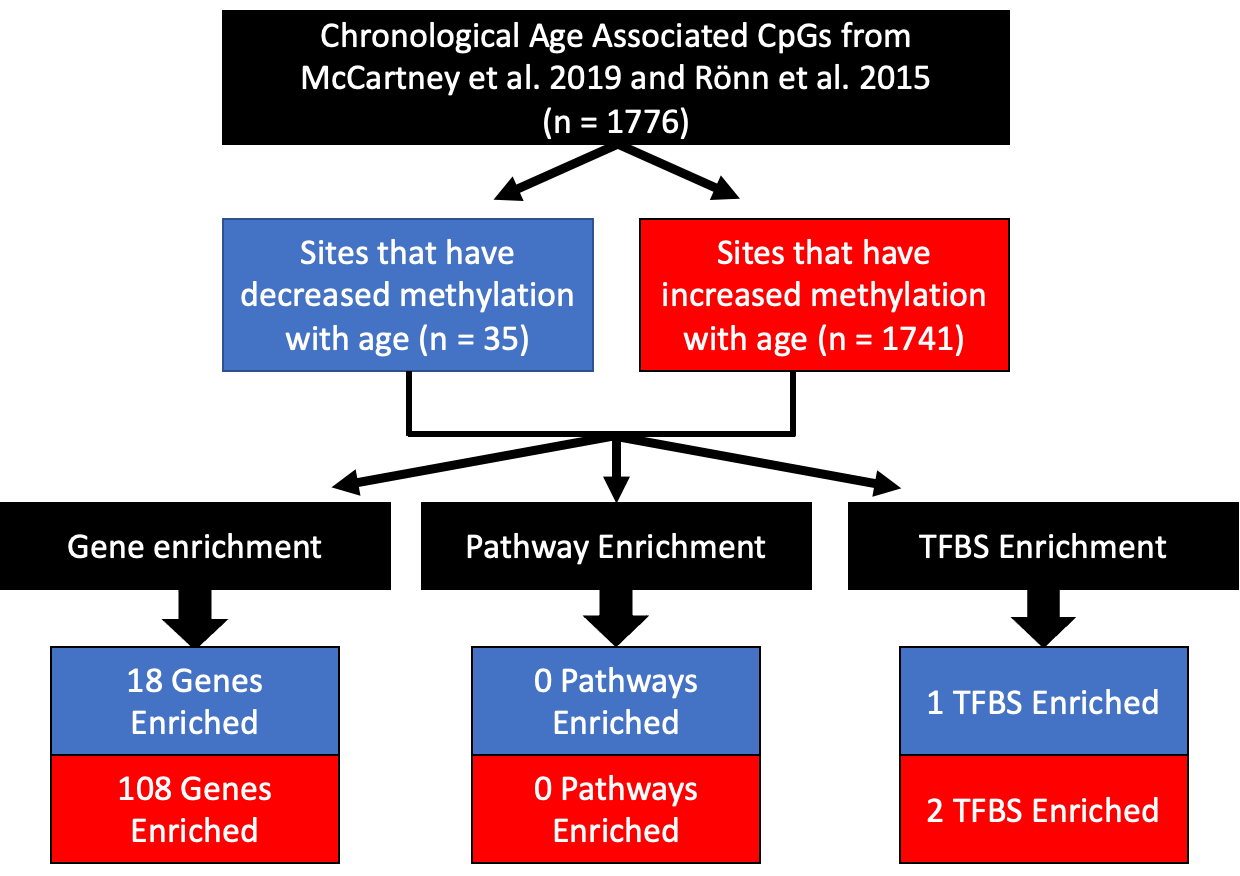
